# Differential responses in the core, active site and peripheral regions of cytochrome c peroxidase to extreme pressure and temperature

**DOI:** 10.1101/2024.07.24.604936

**Authors:** Rebecca K. Zawistowski, Brian R. Crane

**Affiliations:** Department of Chemistry and Chemical Biology, Cornell University, 122 Baker Laboratory Ithaca, NY 14853, USA

**Keywords:** high-pressure crystallography, protein packing, protein folding, side-chain dynamics, solvation, heme-ligation

## Abstract

In consideration of life in extreme environments, the effects of hydrostatic pressure on proteins at the atomic level have drawn substantial interest. Large deviations of temperature and pressure from ambient conditions can shift the free energy landscape of proteins to reveal otherwise lowly populated structural states and even promote unfolding. We report the crystal structure of the heme-containing peroxidase, cytochrome c peroxidase (CcP) at 1.5 and 3.0 kbar and make comparisons to structures determined at 1.0 bar and cryo-temperatures (100 K). Compressibility plateaus after 1.5 kbar and pressure produces anisotropic changes in CcP. CcP responds to pressure with volume declines at the periphery of the protein where B-factors are relatively high but maintains nearly intransient core structure and active site channels. Compression at the surface affects neither alternate side-chain conformers nor B-factors. Thus, packing in the core, which resembles a crystalline solid, limits motion and protects the active site, whereas looser packing at the surface preserves side-chain dynamics. Changes in active-site solvation and heme ligation reveal pressure sensitivity to protein-ligand interactions and reveal a potential docking site for the substrate peroxide.

## Introduction

The properties of proteins depend highly on temperature and pressure [1–5] **^1^**. Higher temperature will generally favor structural states with higher conformational entropy [6–10]. In contrast, higher pressure will favor states that occupy smaller volumes [7,9,11–13]. In both cases, shifting the energy landscape of both proteins and nucleic acids may reveal states that are normally not well populated, yet important for function as key intermediates in reactions or as conformations stabilized by binding partners [14–17]. Nonetheless, determining crystal structures under extreme conditions is challenging. Although there are several deposited structures at high-temperature, few structures have been determined at high-pressures [18].

The effects of high-pressure (HP) on globular proteins are generally governed by the free energy relationship between pressure and volume: with increasing pressure, systems compress to states that occupy less volume. Due to the imperfect packing of globular proteins, extended, unfolded states have decreased molecular volumes because of the loss of cavities and tunnels. Smaller protein volumes can also result from cavity compression, isotropic thermal (B)-factor depression, hydrogen-bond contraction, and loss of alternate side-chain conformations [16,18–20]. Thus, pressure perturbations allow one to probe conformations that are not normally accessible via conventional crystallographic techniques.

The advent of high-pressure crystallographic instruments and techniques such the diamond anvil cell (DAC), has allowed access to pressure-perturbed states at ambient temperature [18,21,22]. For example, in 2001 Fourme and coworkers utilized a DAC to pressurize hen egg white lysozyme (HEWL) and bovine erythrocyte Cu, Zn superoxide dismutase crystals under 10 and 9.0 kbar respectively, and reported the structure of HEWL at 6.9 kbar to high-resolution [21]. These results confirmed previous high-pressure data on HEWL from Kundrots and Richards collected in 1987 that demonstrated a general resistance of the global structure to pressure-induced changes [19]. Since then, a number of high-pressure structures have been determined. For example, the structure of sperm whale myoglobin under 1.5 and 2.0 kbar of pressure revealed conformational substates similar to those observed at low pH [23]. Crystals of macromolecular assembly cowpea mosaic virus (CpMV) showed an increase in diffraction quality under higher pressures [24]. Other recent HP structures include those of ribonuclease A, β-lactoglobulin, and insulin [25–27]. Recently, the HP crystallographic structure of the Ras oncogene protein at 5.0 kbar revealed otherwise “hidden” conformational states that allow for inhibitor binding [14,15].

However, determining crystallographic structures from a DAC is not without challenges. Samples must be mounted in a buffer or medium conducive to high-pressure. Certain crystals may be intolerant to buffer conditions or high-pressure itself. Sufficient penetration of the DAC windows requires intense high-energy synchrotron x-rays, which readily induce crystal damage at the ambient temperatures required for pressurization. Completeness and resolution of the dataset are also often limited by the DAC aperture [18]. Hence, DAC datasets usually require the mounting and pressurization of multiple crystals from which multiple partial datasets are merged.

Cytochrome c peroxidase (CcP) has long stood as a model metalloenzyme to study peroxidase activity and electron-transfer reactions in proteins [28–30]. The protein is readily crystallized and has been used for extensive kinetic and structural analyses [31,32]. CcP catalyzes the reduction of hydrogen peroxide to water with a heme cofactor that is subsequently reduced by cytochrome c [28]. CcP has structurally complex architecture that contains a sizeable channel at its core for exchange of substrates and products to the active center. In addition, networks of hydrogen bonds and van der Waals contacts facilitate long-range electron-transfer from the heme to the protein surface [33,34]. Given the potential of pressure to drastically alter macromolecules and cellular membranes, it is remarkable that extreme environments, such as the deep sea, sustain life [35–37]. How pressure may perturb physical properties in CcP has general relevance for life under extreme conditions and is largely unexplored [24,35,38–41].

In this work, we describe the structure of CcP under two high-pressure conditions and compare those structures to those collected at low and room temperature, and at ambient pressure. Surprisingly, CcP is impervious to large scale changes even at pressures three times those found in the deepest ocean on earth (∼1.0 kbar) [38]. However, the interior of the CcP structure varies less when compared to the peripheral regions, which do exhibit some compression.

Importantly, channels for substrate access are preserved at all pressures and the conformation of the heme cofactor is unperturbed. Surprisingly, the solvation and ligation environment of the heme is pressure dependent. These observations support the idea that globular proteins are efficiently packed in their hydrophobic cores but less so near their surfaces and that pressure may influence the interaction of proteins with solvent and ligand.

## Results

### High-pressure effects on diffraction quality

CcP was crystallized within 24 hrs of data collection to minimize crystal degradation. The crystals formed were red, long and rod-shaped. Diffraction datasets were collected at ambient pressure and cryogenic temperature (AP-Cryo-CcP) and ambient pressure and room temperature (AP-RT-CcP), and at room temperature under two high-pressure conditions, 1.5 kbar (1.5HP-RT-CcP) & 3.0 kbar (3.0HP-RT-CcP), respectively. All of the crystals have the same space group (P2_1_2_1_2_1_) with similar unit cell dimensions. The AP-Cryo-CcP crystals deviate the most from the others in unit-cell lengths, for which *b* and *c* are nearly 2- and 1-Å smaller, respectively, than for the AP-RT-CcP crystals. The unit-cell dimensions for all principal axes in 1.5 kbar structure decrease slightly, whereas the 3.0 kbar structure only has a slight decrease in dimension *a* compared to AP-RT-CcP (**Table 1**). Changes in unit cell parameters are common for cryogenically cooled and high-pressure macromolecular crystals [18,19].

**Table 1.** Data collection and refinement statistics. Values in parentheses are for the outer shell.

|  | AP-RT-CcP | AP-Cryo-CcP | 1.5HP-RT-CcP | 3.0HP-RT-CcP |
| --- | --- | --- | --- | --- |
| PDBid | **9C8L** | **9C8M** | **9C8O** | **9C8P** |
| Pressure (kbar) | 0.001 | 0.001 | 1.5 | 3.0 |
| Resolution range (Å) | 34.92 - 1.54<br>(1.57 - 1.54) | 41.34 - 1.78 (1.81<br>- 1.78) | 26.88 - 2.31 (2.37<br>- 2.31) | 27.49 - 2.31 (2.37<br>- 2.31) |
| Space group | P2<sub>1</sub>2<sub>1</sub>2<sub>1</sub> | P2<sub>1</sub>2<sub>1</sub>2<sub>1</sub> | P2<sub>1</sub>2<sub>1</sub>2<sub>1</sub> | P2<sub>1</sub>2<sub>1</sub>2<sub>1</sub> |
| a, b, c (Å) | 45.14 74.38<br>101.49 | 44.61 72.89<br>100.38 | 44.73 74.05<br>100.89 | 44.71 74.25<br>101.31 |
| α, β, γ (°) | 90 90 90 | 90 90 90 | 90 90 90 | 90 90 90 |
| Total reflections | 343689 | 843312 | 95613 | 117067 |
| Unique reflections | 50414 (2323) | 31873 (1516) | 12171 (881) | 14622 (1096) |
| Completeness (%) | 98.10 (91.85) | 99.52 (97.74) | 79.60 (75.82) | 95.02 (93.36) |
| Redundancy | 3.2 | 26.4 | 5.2 | 5.5 |
| Wilson B-factor (Å<sup>2</sup>) | 21.8 | 24.7 | 35.4 | 26.2 |
| R<sub>merge</sub> (%) | 9.2 (74.7) | 10.6 (90.9) | 23.7 (78.9) | 18.1 (65.4) |
| R<sub>meas</sub> (%) | 10.0 (83.2) | 10.9 (93.6) | 26.2 (87.2) | 20.0 (72.5) |
| R<sub>pim</sub> (%) | 3.9 (35.8) | 2.2 (21.8) | 10.7 (36.3) | 8.2 (29.8) |
| CC<sub>1/2</sub> (%) | 98.7 (60.1) | 100.0 (91.0) | 96.1 (75.0) | 97.3 (65.1) |
| CC* (%) | 99.7 (86.7) | 100.0 (97.6) | 99.0 (92.6) | 99.3 (88.8) |
| Mean I/σ (I) | 20.8 (0.9) | 58.2 (3.7) | 6.7 (1.8) | 7.2 (2.0) |
| Reflections used in refinement | 50414 (2323) | 31873 (1516) | 12171 (881) | 14622 (1096) |
| Reflections used for R-free | 2000 (92) | 2000 (95) | 1217 (88) | 1461 (109) |
| R-work (%) | 15.5 (36.5) | 17.3 (27.0) | 19.3 (23.1) | 21.7 (27.4) |
| R-free (%) | 18.0 (38.1) | 21.3 (31.6) | 25.0 (29.2) | 27.3 (33.8) |
| Number of non-hydrogen atoms | 2643 | 2749 | 2607 | 2610 |
| macromolecules | 2398 | 2387 | 2353 | 2346 |
| ligands | 43 | 43 | 45 | 43 |
| solvent | 202 | 319 | 209 | 221 |
| Protein residues | 296 | 295 | 295 | 295 |
| RMS(bonds) | 0.011 | 0.006 | 0.002 | 0.002 |
| RMS(angles) | 1.1 | 0.86 | 0.49 | 0.53 |
| Ramachandran favored (%) | 99.7 | 98.6 | 98.3 | 98.3 |
| Ramachandran allowed (%) | 0.34 | 1.37 | 1.71 | 1.37 |
| Ramachandran outliers (%) | 0 | 0 | 0 | 0.34 |
| Rotamer outliers (%) | 0.39 | 1.2 | 0 | 1.26 |
| Clashscore | 2.11 | 1.06 | 3.47 | 2.84 |
| Average B-factor | 26.0 | 26.9 | 37.4 | 28.5 |
| macromolecules | 25.2 | 25.7 | 37.0 | 28.4 |
| ligands | 15.9 | 17.1 | 28.6 | 21.1 |
| solvent | 37.2 | 36.6 | 43.1 | 31.0 |
| Average H-bond distances (Å) | 3.02 ± 0.15 | 3.02 ± 0.15 | 3.04 ± 0.18 | 3.05 ± 0.17 |

Notably, the high-pressure structures are of lower resolution compared to AP-RT-CcP and AP-Cryo-CcP (**Table 1**). Loss of resolution at HP is due at least in part to the detrimental effects of NVH oil immersion. Crystals under high pressure were also measured in mother liquor solution in attempts to collect higher resolution data. However, diffraction data from such crystals were inconsistent and could not be merged to produce a complete dataset.

Despite reported decreases in hydrogen bond distances in high-pressure protein structures when compared to the same proteins at ambient pressure, we do not observe significant changes in the average hydrogen bond distance when the proteins are under pressure (**Table 1**) [19,20]. Instead, average distances are maintained at around 3.0 Å regardless of temperature or pressure conditions.

### Differences in compression at two different pressures

Visual inspection of C_α_ backbone superpositions for the different pressure conditions show little deviation among the structures (**Figure 1**). Difference distance matrices of the average atomic distances of each individual residue vs every other residue indicate a modest overall compression of the structures at the higher pressures (**Figure 2a & b**). The average difference distance for each residue relative to all other residues was mapped onto to a ribbon representation of CcP to reveal regions of the molecule that the rest of the protein has moved toward or away from, on average. The 1.5 kbar structure (1.5HP-RT-CcP) is globally compressed compared to that at ambient pressure (**Figure 2a**). In contrast, the 3.0 kbar structure (3.0HP-RT-CcP) shows compression compared to the 1.0 bar structure (AP-RT-CcP), but the differences are more anisotropic than with 1.5HP-RT-CcP, with some regions showing greater compression than others (**Figure 2b**).

**Figure 1.**
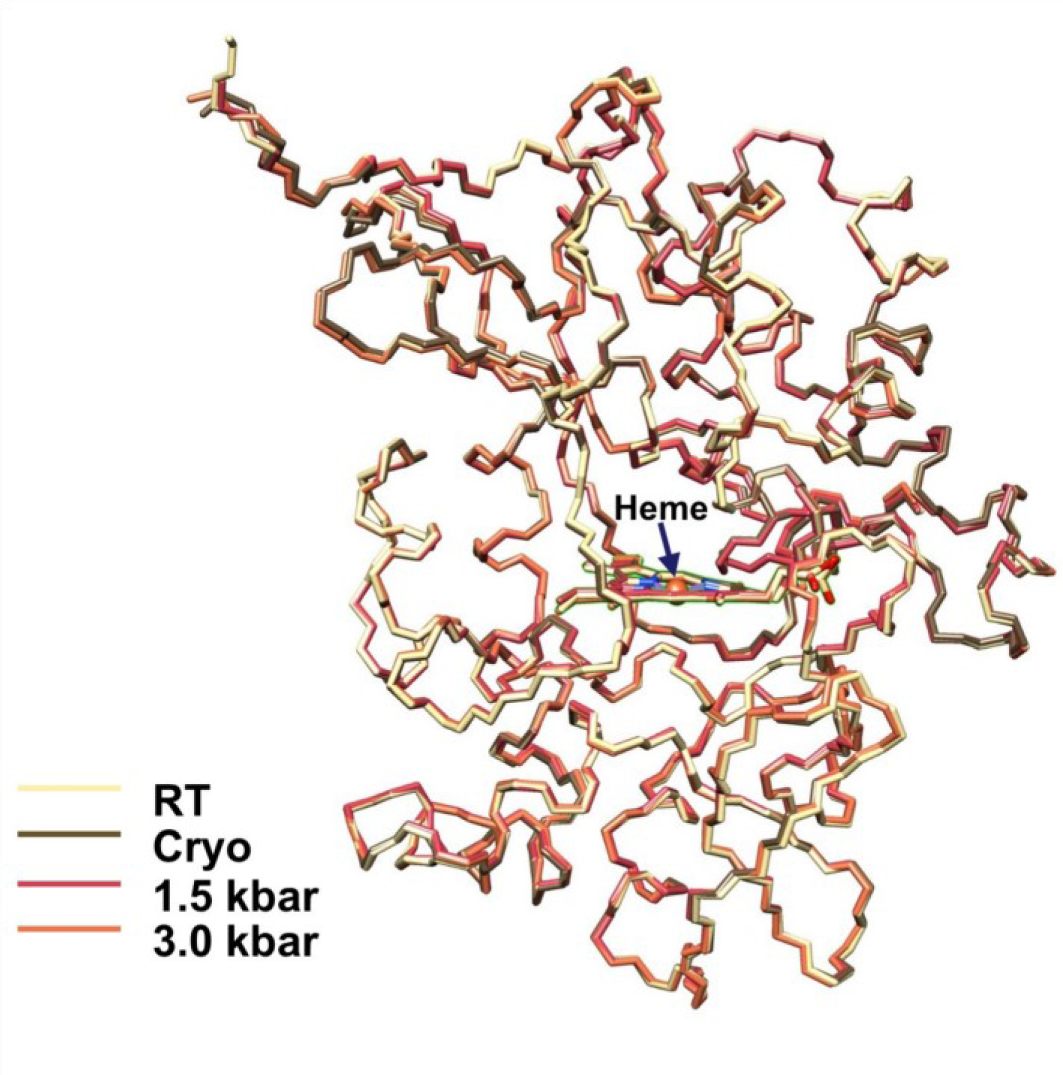
Superpositions of AP-RT-CcP (PDB Code **<u>9C8L)</u>**, AP-Cryo-CcP (**<u>9C8M)</u>**, 1.5HP-RT-CcP (**<u>9C8O)</u>**, and 3.0HP-RT-CcP (**<u>9C8P)</u>**. Global comparison of C_α_ positions reveals that neither large changes in temperature nor pressure have affected the overall conformation of the protein.

**Figure 2.**
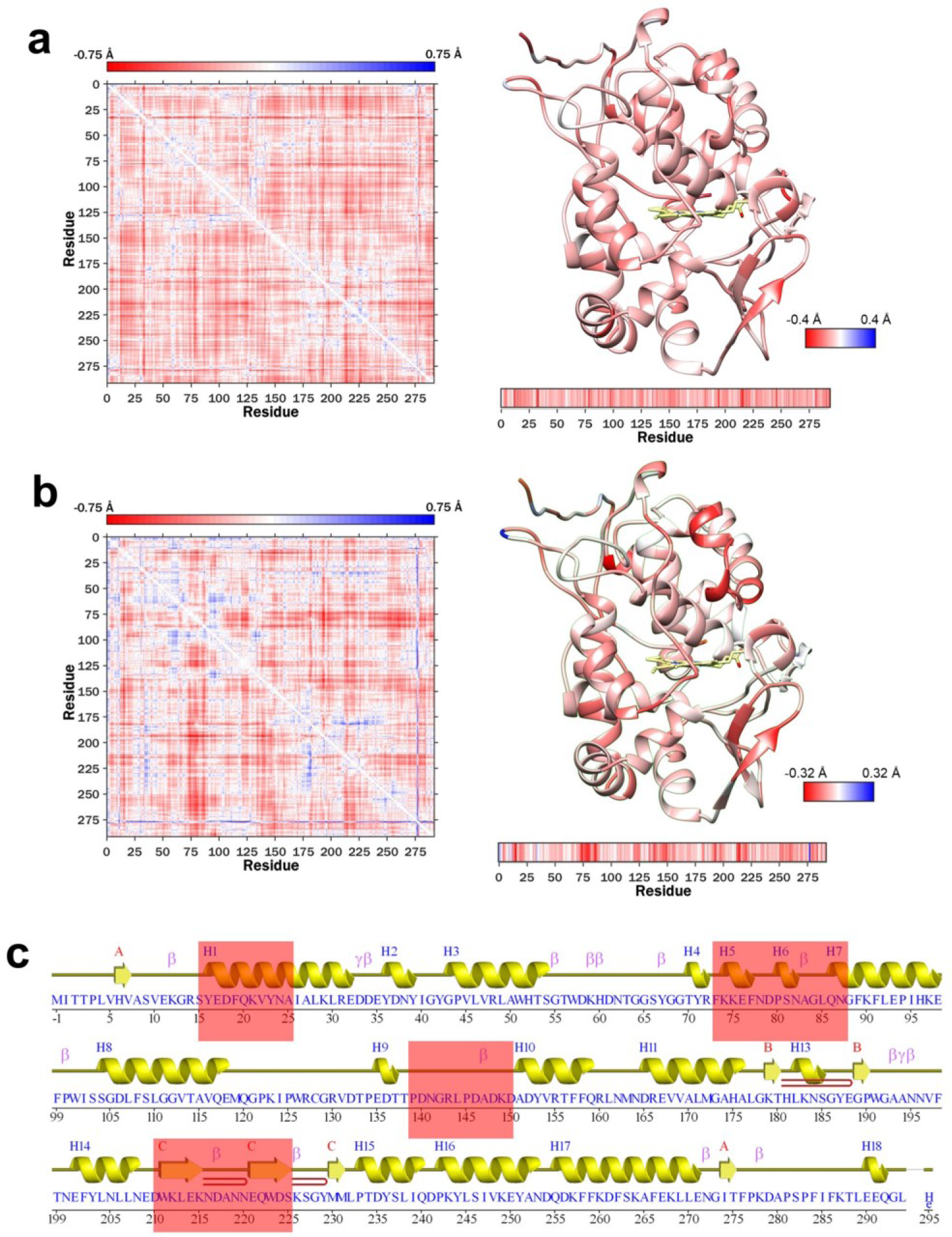
Difference distance maps for 1.5HP-RT-CcP – AP-RT-CcP (**a**) and 3.0HP-RT-CcP – AP-RTCcP (**b**). Overall, the high-pressure structures contract compared to the RT structure. The 3.0 kbar CcP structure exhibits specific regions of compression whereas the 1.5 kbar structure undergoes a more isotropic compression. (**c**) Mapping of regions of compression in 3.0HP-RT-CcP to the secondary structure elements.

In 3.0HP-RT-CcP the regions of greatest change map to distinct secondary structure features, particularly α-helices and β-strands on the periphery of the protein (**Figure 2b & c**), whereas perturbations within the protein core are very small compared to the ambient pressure structure. This pattern of contraction is similar to that seen when comparing AP-Cryo-CcP to AP-RT-CcP, wherein the major distance changes appear primarily on the outer parts of the protein and the central regions remain mostly fixed.

The compressibilities of the HP structures (-1/V x ∂V/∂p) were calculated from the differences in molecular volumes (V_MS_) of CcP between the two HP conditions and ambient pressure. The compressibility of CcP is nonlinear with the ambient to 1.5 kbar change generating greater compressibility (14.0 Mbar^-1^) than the ambient to 3.0 kbar change (5.74 Mbar^-1^). The compressibility between 1.5 and 3.0 kbar was calculated to be -2.61 Mbar^-1^. The negative value owes to a modest volume increase between the two high pressure conditions (from 45007 Å^3^ to 45183 Å^3^) that is likely within the error of the structural modeling. Nevertheless, the decrease in compressibility between 1.5 and 3.0 kbar reflects an initial compaction of the protein to 1.5 kbar and then little additional compression with increasing pressure.

### Differences in heme active-site features at different pressures

Although the heme cofactor does not show any perturbations among the structures, solvation of the distal heme pocket does differ considerably among them. Four ordered water molecules, including one that coordinates the ferric heme iron, are most readily discerned in AP-Cryo-CcP. The heme-coordinating solvent molecule has considerably more delocalized density at room temperature than in the cryo structure. Notably, the 1.5HP-RT-CcP diffraction data produces oblong electron density in the F_o_-F_c_ omit map that is well-modeled as a diatomic molecule (for example H_2_O_2_, O_2_ ^2-^, O_2_ ^-^ or O_2_) (**Figure 3a-d**). Attempts to attribute this density to two water molecules produced contacts inside of van der Waals distances and increased the R_Free_ statistics (**Figure 3e**). Furthermore, the heme-bound water found in the other structures is not present in 1.5HP-RT-CcP, likely owing to unfavorable van der Waals contacts with the diatomic species. Unexpectedly, this diatomic-shaped electron density is not evident in the 3.0HP-RT-CcP data, which rather shows a water molecule pattern more similar to that of the ambient pressure structures, including density for the heme-bound water molecule. Similar to 1.5HP-RT-CcP, the ordered water molecules in the heme pocket of 3.0HP-RT-CcP are not as well defined as in either of the ambient pressure structures (**Figure 3d**).

**Figure 3.**
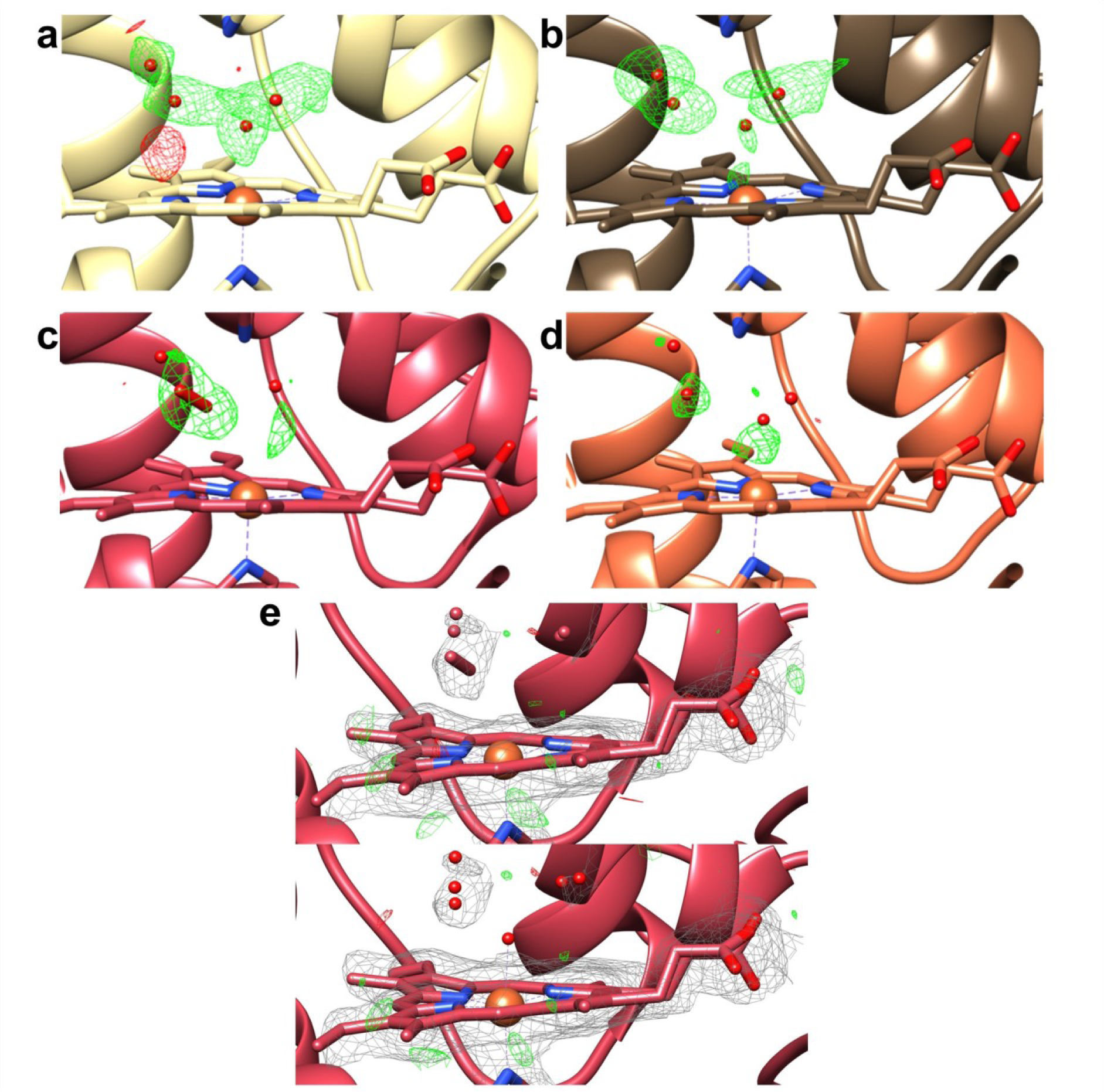
F_o_-F_c_ omit maps contoured to 2.5 σ (green positive, red negative) in the heme bound active site in AP-RT-CcP (**a**), AP-Cryo-CcP (**b**), 1.5HP-RT-CcP (**c**), and 3.0HP-RT-CcP (**d**). The 1.5 kbar structure exhibits density in the map that fits well to a diatomic molecule (possibly O_2_^2-^ or O_2_) that is not found in the 3.0 kbar structure. (**e**) Refinement of 1.5HP-RT-CcP with two water molecules instead of a diatomic O_2_^x-^ molecule (upper panel) does not fit the 2F_o_-F_c_ map (grey, contoured to 1 σ) as well as a diatomic O_2_^2-^ molecule (lower panel) and increases the R_Free_ statistics during refinement. Unlike the other structures, there is no density for a heme-coordinating water molecule.

### Tunnel and cavity volume decline correlates with increasing pressure

The MOLE2.0 [42] cavity search software was applied to analyze cavity and tunnel changes within the structures (**Figure 4a**). The interior threshold probe radius used for the cavity search (0.7 Å) was less than that of a solvent molecule in order to detect small changes in the spaces within the structures. The tunnels that the algorithm identified differ depending on pressure. Many of the tunnels found at ambient pressure disappear at 3.0 kbar. The location of the tunnel losses correlate with the compression regions identified in the distance difference matrix between 3.0HP-RT-CcP and AP-RT-CcP (**Figure 4b**).

**Figure 4.**
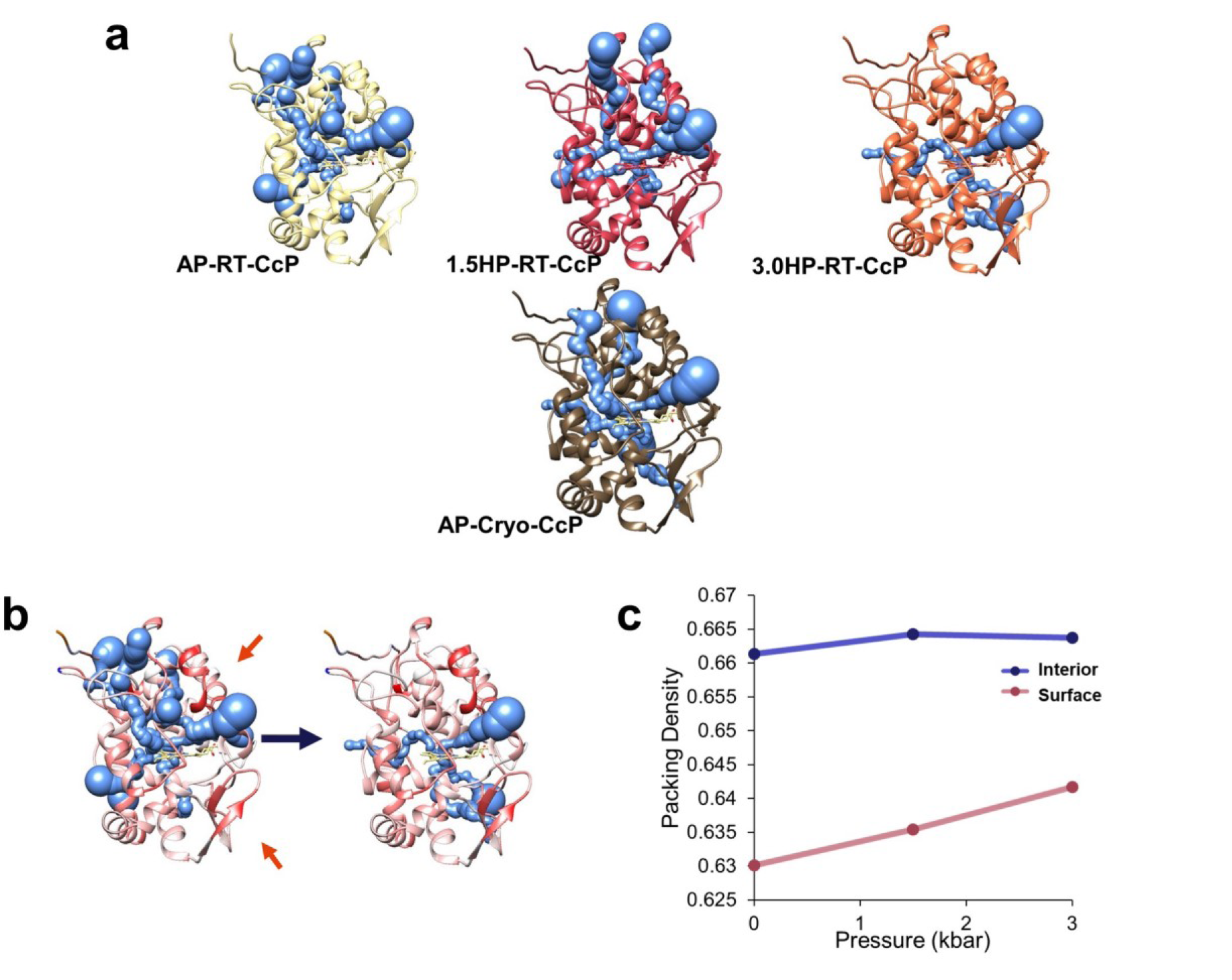
Cavity analysis for CcP structures under temperature and pressure conditions. (**a**) Tunnels for each of the ambient pressure and high-pressure models calculated with the MOLE2.0 cavity search algorithm (see Methods). (**b**) Tunnels for RT-AP-CcP and 3.0HP-RT-CcP superimposed on difference distance model from Figure 2B (more red regions represent greater compression). Tunnels in the 3.0 kbar structure match closely with mapped compression regions in the structures (noted by the red arrows). (**c**) Interior and surface packing densities with respect to pressure. The interior packing densities are larger in all of the structures than the surface packing densities, which increase with increasing pressure.

Packing densities for both the interior and surface of the protein were calculated for each of the structures. The interiors of the CcP structures pack more tightly than at the surfaces, a finding that agrees with previous assessments of packing densities in proteins [43]. Notably, the packing density of the interior is not greatly perturbed with increasing pressure, however; the peripheral regions that are more close to the protein surface, exhibit an increase in packing density with higher pressures (**Figure 4c**).

### Static disorder can dominate positional uncertainty in CcP crystals

The average isotropic B-factor per residue (B_iso_) was highly consistent across the structures when they were refined to highest common resolution (**Figure 5a**). Nonetheless, 1.5HP-RT-CcP initially gave an apparent increase in B_iso_ across all residues. However, excluding one of the four crystals that contributed to the overall dataset abrogated the effect (**Figure 5b**). Thus, crystal heterogeneity stemming largely from one outlier crystal was responsible for the apparent change in B-factors for the 1.5 kbar structure. Surprisingly, there are no major differences in B-factors between the two structures collected at different temperatures. Hence, static disorder caused by differences in corresponding atomic positions across unit cells likely contributes differentially to the atomic displacement parameters in these structures. Perturbations to the lattice induced by cryocooling may increase the relative average B-factors and compensate for reduced fluctuations with temperature.

**Figure 5.**
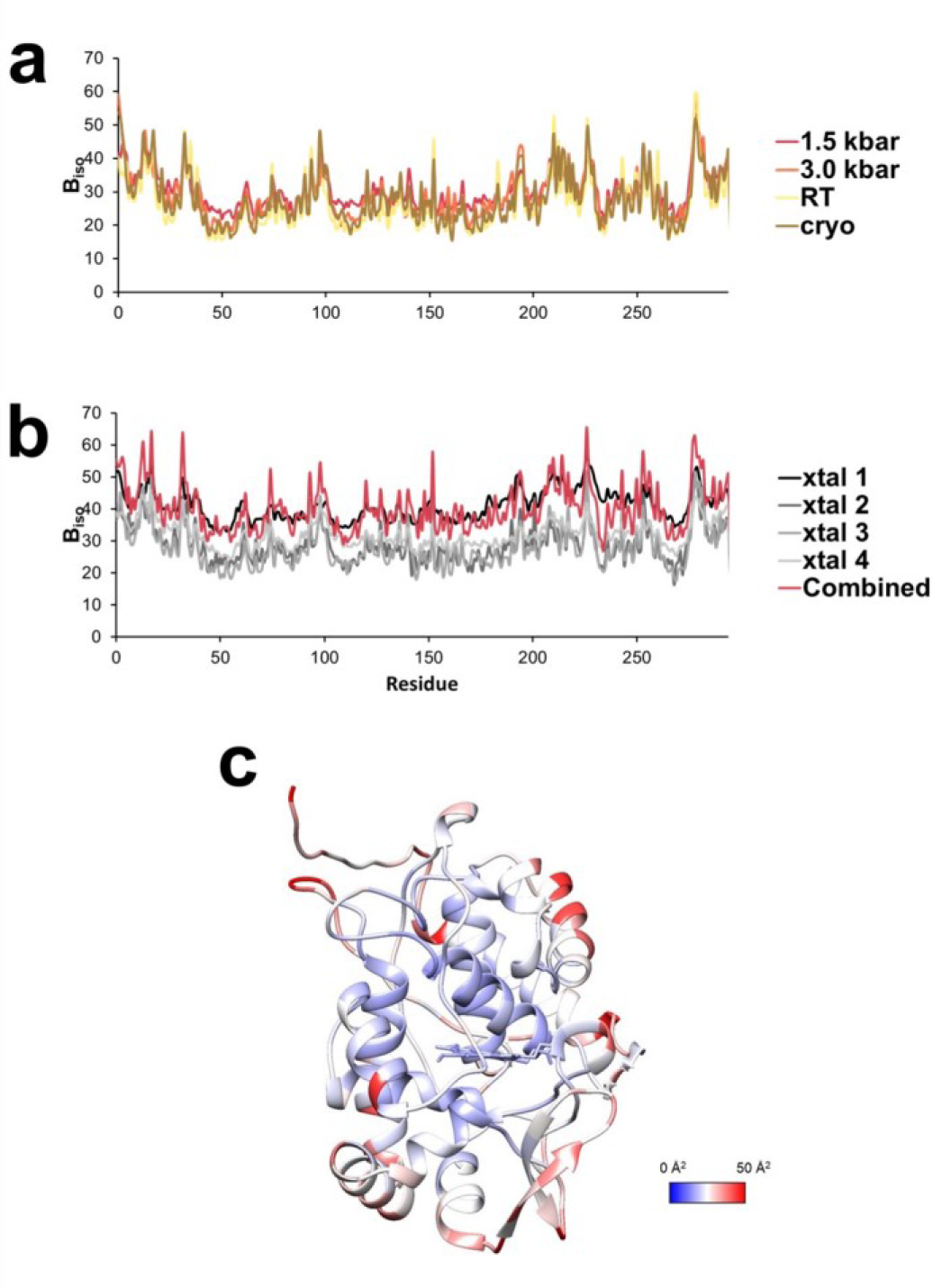
(**a**) Average isotopic B-factors for all models when refined to 2.3 Å resolution. There are no major changes in B-factors across the protein when comparing different conditions. Notably, the 1.5HP-RT-CcP, showed an overall baseline increase which is largely due to the merging of multiple crystals (**b**). Crystal (xtal) 1 gives higher overall B-factors than the others and was thus omitted in the final model and when comparing B-factors between models. (**c**) Average B-factors mapped onto the AP-RT-CcP structure. B-factors (values indicated by the color legend) are lower in the interior and near the active site core than at the periphery.

When mapping average B_iso_ onto the structures, a clear differentiation becomes apparent between the interior of the model and the surface regardless of temperature or pressure condition (**Figure 5c**). The interior of the protein has relatively low B_iso_ compared to the exterior, with the lowest B_iso_ values being found in the heme pocket.

### Different pressures do not favor specific side-chain conformers

The multiconformer search algorithm qFit [44] was used to identify possible multiple conformers of residues in each of the structures. For AP-Cryo-CcP, 1.5HP-RT-CcP, and 3.0HP-RT-CcP, qFit identified several residues with multiple conformers; however, upon inspection of difference electron density maps, these conformers could not be unambiguously established above the noise levels of the maps. However, the 1.54 Å resolution diffraction data of AP-RT-CcP revealed multiple conformers for residues with clear positive density in the F_o_-F_c_ map (**Figure 6a,b**). Furthermore, inclusion of these multiple conformers for Thr156 and Ser237 in the models reduced the R_Free_ statistics of AP-RT-CcP. Positive difference density indicative of the multiple conformations of these residues can be seen in the high-pressure structures as well, and they were thus included in the corresponding refined models (**Figure 6c**). Thus, the multiple conformers do not appear specific for a given temperature or pressure.

**Figure 6.**
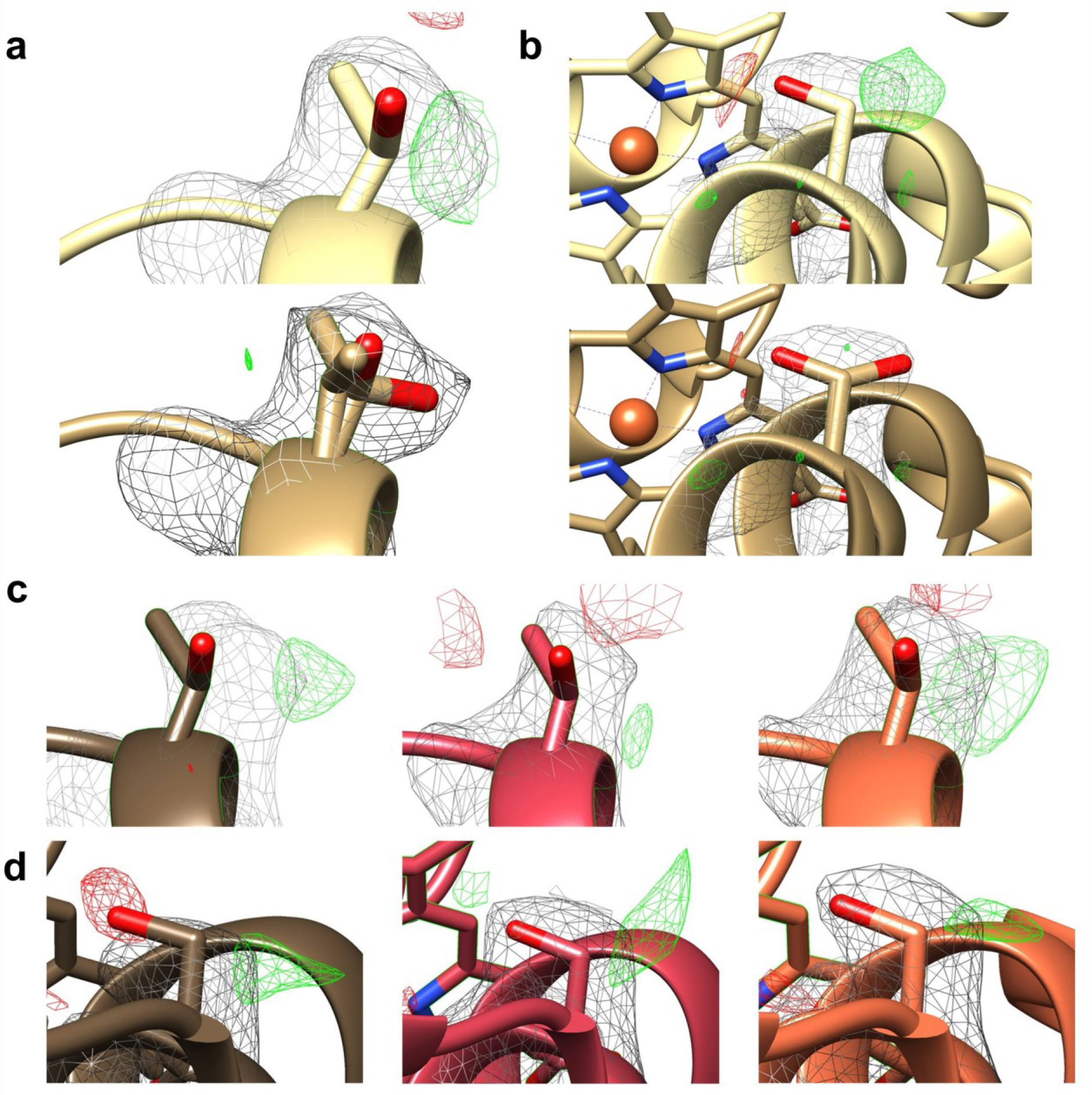
Refinement of multiple conformer residues identified by qFit in AP-RT-CcP vs single conformer model refined to same densities. 2F_o_ - F_c_ maps all contoured to 1 σ (grey). Positive electron density is observed in the difference map (contoured to 2 σ, green positive, red negative) when refined to a single conformer for residues (**a**) Thr156 and (**b**) Ser237. F_o_ - F_c_ maps (contoured to 2 σ) of (**c**) Thr156 and (**d**) Ser237 for AP-Cryo-CcP (brown), 1.5HP-RT-CcP (red), and 3.0HP-RT-CcP (orange) show similar patterns of positive (green) and negative (red) difference electron density indicative of multiple conformations.

## Discussion

The effects of pressure on chemical equilibrium is related to changes volume [6,18,38]:

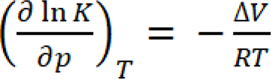

Where *K* is the equilibrium constant, *p* is pressure, *T* is temperature, and ΔV is the change in volume. As such, increasing pressure will shift the ensemble towards states which occupy smaller volumes. We observe that the CcP crystal structure is largely invariant to both temperature and pressure perturbations. These structures show that CcP in crystals maintains a single, tightly clustered conformational ensemble independent of higher temperatures and pressures.

Increased pressure usually favors less-globular states over fully folded states, which tend to have cavities that increase overall volume [18,20]. However, for CcP the native fold stays intact at high pressure with only a slight decrease in cavity and tunnel volumes. The lack of large changes may derive from well-packed, interior solvent acting as a pressure medium in the crystals, as has been noted previously for hen egg-white lysozyme crystals [22,45]. In the case of the Ras protein, where pressure induced conformational changes have been observed, the protein under study is a signal transduction protein, which must readily change conformation to deliver a signaling function [14,15]. An enzyme like CcP, whose catalytic reactivity is largely determined by its cofactor-bound active site, may not have energetically low-lying more extended conformational states available.

The low compressibility of proteins stems from complimentary packing of the protein residues and ordered solvent [46]. Compressibility measurements indicate that CcP contracts more between ambient and 1.5 kbar than between 1.5 and 3.0 kbar, which would be expected for a structure that is intrinsically well packed under most conditions. Notably, the protein largely resists further compression at pressures beyond that found on the earth’s surface. It has been suggested that for globular proteins, packing in the interior of the protein is greater than near the surface [43]. The trend of pressure increasing packing density near the surface of the CcP structures verifies this finding. Furthermore, the anisotropic compression in 3.0HP-RT-CcP structure closely correlates with cavity and tunnel volume decreases. Thus, the well-packed core of CcP resists compression and restricts pressure-induced structural changes. In contrast, protein structure near the surface is more malleable and can decrease volume to minimize free energy. Packing densities within the core of globular proteins are similar to those of crystalline solids [43,47,48]. As such, we would expect similar pressure-induced effects for other globular proteins at the periphery where packing density is relatively lower than the core.

Another way that protein structure can potentially decrease volume is through the compression of hydrogen bonds [39,40]. NMR and crystallography studies indicate that high-pressure strengthens hydrogen bonds in proteins [24,38–41]. Changes in overall hydrogen bonding distances within CcP cannot be resolved at these resolutions. Rather, the structure of CcP compensates for pressure by decreasing cavity and tunnel sizes in areas where the residues are loosely packed and not through the shortening of hydrogen bonds.

Side-chain conformers favored by extreme conditions have been observed for many higher-temperature (> 20 °C) and some high-pressure crystal structures [16,49–51]. In the case of CcP protein crystals, however, we observe the same set of conformers, regardless of temperature and pressure. For example, Thr156 and Ser237 display the same alternate side-chain conformers in all structures. These conformations likely have similar energy and thus one is not necessarily preferred by lower temperature. It is unlikely that the conformers are trapped in their ambient temperature positions upon flash-cooling because crystals cool to cryogenic temperatures in seconds [52,53] and side chain motions occur on the nano-second-to-micro-second time scale [54]. It may be no accident that both Thr156 and Ser237 are near the protein surface with Thr156 solvent-exposed. Although, pressure-induced volume reduction could select for one conformer, either conformer may be equally accommodated by a volume reduction with pressure thereby “locking-in” one or the other. Even as surface side chains become more tightly packed, they can maintain fluctuations through correlated interactions and motions [55]. Indeed, the hydrophobic core of globular proteins hinders side-chain flexibility, yet in these more densely packed regions, concerted motions dominate [56,57]. Although packing density increases at higher pressures on the surface, these densities are still relatively small (< 65%) and less than the packing at the core. Packing densities do correlate with B-factors. For globular proteins, B_iso_ values tend to be smaller at or near the active site of the protein indicating greater structural homogeneity [58,59]. In CcP, interior residues, near the active site, generally have lower B_iso_ values than at the protein surface. Furthermore, the areas with higher B_iso_ values also correspond to regions that compress more under 3.0 kbar when compared to ambient pressure. Thus, the structures are more restricted in the interior than at the surface regions and the protein cores are more impervious to change [28,30].

B-factors reflect both dynamic and static disorder, with the latter arising from different atomic positions in the crystal relative to the respective unit-cell origins and axes [60,61]. Such variability can derive from intrinsic crystal packing, conformational heterogeneity, or be induced by cryo-cooling and radiation damage [62]. Rigid body vibrations of the molecules across unit cells within the lattice can also contribute to the individual B-factors, without reflecting relative motions within the proteins themselves [63]. Thermal motion of the atoms is influenced by protein conformation and globular structure and gives rise to a larger average B_iso_ values in higher temperature structures [58,64,65]. Global analysis of structures in the protein data bank and systematic studies on individual proteins show that static disorder dominates B-factors at low to medium resolution and at low temperature (> 80%) [63,66,67]. Only in high-resolution non-cryo structures does the major contribution of the B-factor derive from thermal vibrations dependent on the protein structure (> 60% at > 1.2 Å resolution). Moreover, static disorder increases at cryo-temperatures to compensate for the decline in thermal motion such that there is very little statistical difference between the B-factors of proteins whose data was collected at 100 vs 300 K [66]. Some high-pressure structures exhibit lower B-factors than their ambient temperature counterparts, which may be indicative of restricted motion upon compression [19]. However, for the CcP crystals neither temperature nor pressure influence B_iso_ values compared on a per residue basis or as overall Wilson B-factors (Table 1). In fact, the highest overall B-factor of 1.5HP-RT-CcP may derive from crystal heterogeneity because removing one crystal from the dataset greatly influenced the overall B-factor. It is intriguing that B_iso_ values do not change significantly with pressure for CcP; even with an increase in packing and compression at the protein surface, the residues themselves do not have lower atomic displacement parameters. The insensitivity of B-factors to these modest changes in compression suggests that the fluctuations governing the B_iso_ values are too small in amplitude to be affected by volume the losses, and that unit cell static disorder generally dominates the B-factors.

The most substantial pressure effects involve the ordered solvent of the heme pocket. Ordered water molecules are well-defined in AP-Cryo-CcP, have a similar pattern in AP-RT-CcP and are less well discerned in the high-pressure structures. However, the largest changes to solvation of the heme pocket are at midrange pressures. In 1.5HP-RT-CcP, electron density to the side of the heme iron is fit best by a diatomic molecule. CcP reacts with hydrogen peroxide as a substrate, and hydrogen peroxide could be present in the PEG-containing buffer or generated by the reducing x-ray beam at room temperature [68,69]. Furthermore, hydrogen peroxide that is not ligated by metal ions has previously been visualized in the active sites of proteins [70]. Nonetheless, it would be unusual for hydrogen peroxide to localize beside the heme iron and not react unless the heme iron is also reduced. Photoreduction of heme via x-ray radiation during the course of crystallographic measurements has been observed and often convolutes structural information, such as metal ligation, around the cofactor [71]. CcP crystals appeared visually to change in color over the course of x-ray diffraction measurements. Though these changes were not measured spectroscopically, it is possible that the iron is being reduced to prevent hydrogen binding. Oxygen is another candidate for the identity of the diatomic species, but seems less likely given its lack of charge and highly dynamic nature. Thus, the density was modelled as peroxide rather than oxygen (the protonation state of peroxide is undetermined and could be H_2_O_2_, HO ^-^ or O_2_ ^2-^). Regardless of the molecular identity, it is striking that the oblong density does not persist in 3.0HP-RT-CcP. This behavior emphasizes how specific pressures may favor specific solvation states. Moreover, the specific conditions of 1.5HP-RT-CcP potentially resolve peroxide in a pre-ligation configuration that reveals a docking site for the substrate within the active center.

### Conclusions

Despite challenges in sample preparation and data collection, high-pressure crystallography is a useful tool for exploring the conformational landscape of proteins as well as their interactions with solvent and ligands. Altering the physical parameters, temperature and pressure of CcP protein crystals does not significantly impact the protein structures. At higher pressures, distinct regions at the periphery of the protein contract, but the core is resistant to compression. This finding verifies that globular proteins are less well-packed on the surface, and that regions with lower packing densities and higher B-factors will undergo the most compression under high-pressure conditions. Although the surface packing densities increase with pressure, we find that specific alternate conformers near the surface are not preferentially favored and that active site channels remain intact. These analyses point to the rigidity and stability of the active-site core in globular proteins as well as provide insight into the ability of proteins to preserve side chain dynamics in tightly packed regions. The most noticeable changes to the CcP active site are at midrange pressures, where altered solvation patterns in the active site suggest that pressure changes could influence interactions between the protein and ligands.

## Materials & Methods

### Protein expression, purification, and crystallization

CcP was expressed and purified as described previously [29,72]. *Escherichia coli* BL21(DE3) cells were transformed with the CcP gene in a ppSUMO vector and grown at 37°C in LB with 50 μg/mL kanamycin. When the OD_600_ reached ∼0.8, cells were induced with 100 μM isopropyl β-D-1-thiogalactopyranoside and expressed at 20 °C overnight. Cells were lysed via sonication and soluble protein was isolated by centrifugation. CcP was purified from lysate using a Ni-NTA column and His-SUMO tags were cleaved with ULP-1 protease. Tags were removed on a Ni-NTA column and CcP was collected in the flow through. The protein was then stirred overnight at 4 °C with 1 molar equivalent of hemin dissolved in 0.1 M NaOH. The reaction was neutralized with acetic acid and centrifuged to remove precipitated heme. Heme containing CcP was then purified via size exclusion chromatography and anion exchange chromatography. The protein sample was concentrated, flash frozen in liquid nitrogen, and stored at -80°C.

Prior to crystallization, Fe CcP was buffer exchanged into filtered Nanopure water and the concentration was diluted to 1 mM. Initial crystal hits were obtained using the Gryphon robot (Art Robbins Instruments). Larger crystals were optimized and grown via vapor diffusion in a 10 μL sitting drop against a reservoir containing 10-25 % polyethylene glycol 550 (MME) and 100 mM 2-(*N*-morpholino)ethanesulfonic acid (pH 6.1-7.1).

### Crystal mounting, data collection, and data processing

Diffraction data were collected at either Brookhaven National Laboratory (BNL) 17-ID-2 FMX beamline on an Eiger 16M detector for ambient pressure, room temperature data or at the Cornell High Energy Synchrotron Source (CHESS) 1D7B2 beamline on an Eiger2 16M detector for high pressure, room temperature and cryogenic data. For high pressure measurements, 1-2 CcP crystals were mounted along with a ruby crystal in a diamond anvil cell (DAC) [21] containing NVH oil. First, a ruby crystal was gently placed into the DAC gasket. NVH oil was then added over the ruby to create a liquid environment for protein crystals and prevent drying. Crystals were then removed from crystallization drops and briefly soaked in oil and then quickly transferred into the DAC gasket. A loop or needle was then used to reorient the crystal in the oil so that as much diffraction data could be collected on the crystal as possible. The DAC was sealed, and compressed with N_2_ gas. Pressures within the DAC were measured by observing wavelength shifts in the ruby fluorescence when excited with 532 nm light. Diffraction data were collected at pressures of 1.5 and 3.0 kbar.

For high-pressure data collection the DAC was rotated 44° about a vertical axis to provide diffraction recording from the full range allowed by the diamond window. Diffraction datasets were indexed, scaled, and merged using HKL-2000 [73]. Scaled sets were then phased via molecular replacement (PDB: 2cyp) and refined using Phaser and phenix.refine in Phenix [74] respectively. Model building was performed with Coot [75].

### Crystal structure comparisons

All four structures were aligned with the MatchMaker function in UCSF Chimera [76] to visualize and compare differences among the structures. Distance difference matrices were calculated using the RR Distance Maps function in UCSF Chimera to compare C_α_ distances from each residue to every other residue within the structures.

Cavity and packing density analyses were performed on each of the structures. MOLE 2.0 [42] was used to calculate cavities within the peptide using a *probe radius* of 5 Å and an *interior threshold* of 0.7 Å. Tunnels were calculated and restricted by a *bottleneck radius* of 0.9 Å. For calculating packing densities, cavities were calculated using an *interior threshold* of 1.4 Å (radius of water). The van der Waals (vdW) and molecular volumes were calculated in MoloVol [77] using the same restrictions (large probe radius: 5 Å, small probe radius: 1.4 Å). The packing densities *P* were then calculated by dividing the vdW volume by the envelope volume:

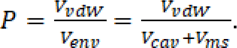

where *V_vdW_* is the vdW volume, *V_cav_* is the cavity (interior or surface) volume and *V_ms_* is the volume of the molecular surface or molecular volume. To compare the packing within the interior core of the protein and the surface, cavities were differentiated as described by Liang and Dill [43], and packing densities were calculated for the interior and surface (*P_int_* and *P_sur_* respectively).

Volume compressibility (β_V_) values were calculated by comparing the change in volume versus the change in pressure:

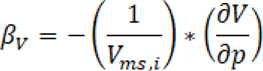

Where *V_ms,i_* is the initial molecular volume at ambient pressure, *∂V* is the relative change in volume from ambient pressure, and *∂p* is the change in pressure.

Multiple side-chain conformers were identified in the ambient pressure, room temperature structures using the multiconformer search algorithm qFit 3 [44]. Electron density around multiconformer residues were then inspected to confirm alternate conformers. Multiple conformer residues were modelled in Coot and B-factors and occupancies were refined in Phenix [74,75].

### Ascension numbers

All coordinates and corresponding structure factors have been deposited to the protein data bank and have the following entries: PDB ID: 9C8L (AP-RT-CcP), PDB ID: 9C8M (AP-Cryo-CcP), PDB ID: 9C8O (1.5HP-RT-CcP), and PDB ID: 9C8P (3.0HP-RT-CcP)

## Abbreviations

DAC: diamond anvil cell
HP: high-pressure
HEWL: hen egg white lysozyme
CpMV: cowpea mosaic virus
CcP: cytochrome c peroxidase
AP-Cryo-CcP: CcP structure at 1.0 bar and cryogenic temperature
AP-RT-CcP: CcP structure at 1.0 bar and room temperature
1.5HP-RT-CcP: CcP structure at 1.5 kbar and room temperature
3.0HP-RT-CcP: CcP structure at 3.0 kbar and room temperature
BNL: Brookhaven National Lab
CHESS: Cornell High Energy Synchrotron Source
vdW: van der Waals
Cav: cavity
MS: molecular surface
Int: interior
Sur: surface

## Acknowledgements

We would like to thank and acknowledge Dr. Zhongwu Wang and Dr. Steve Meisburger at CHESS for their support in data collection. We would also like to thank Dr. Sol Gruner for stimulating discussion of the results.

Secondary structure domain map was constructed using PDBsum on EMBL’s European

Bioinformatics Institute’s webserver.

This work is supported by NSF grant MCB 2129729 to BRC

## Notes

### Competing Interest Statement

The authors have declared no competing interest.

